# Structure of human TRPM8 channel

**DOI:** 10.1101/2022.10.19.512915

**Authors:** Sergii Palchevskyi, Mariusz Czarnocki-Cieciura, Giulio Vistoli, Silvia Gervasoni, Elżbieta Nowak, Andrea R. Beccari, Marcin Nowotny, Carmine Talarico

**Affiliations:** Laboratory of Protein Structure, International Institute of Molecular and Cell Biology, 02-109 Warsaw, Poland; Department of Cell Signaling, Institute of Molecular Biology and Genetics NASU, 03143 Kyiv, Ukraine; Dipartimento di Scienze Farmaceutiche, Università degli Studi di Milano, Via Mangiagalli, 25, I-20133 Milano, Italy; Department of Physics, University of Cagliari, I-09042 Monserrato, Italy; Dompé Farmaceutici SpA, EXSCALATE, Via Tommaso De Amicis, 95, I-80131 Napoli, Italy

## Abstract

**SUMMARY:** TRPM8 is a calcium ion channel that is activated by multiple factors, such as temperature, voltage, pressure, and osmolality. It is a therapeutic target for anticancer drug development, and its modulators can be utilized for several pathological conditions. Here, we present a cryo-electron microscopy structure of a human TRPM8 channel in the closed state that was solved at 2.7 Å resolution. Based on our reconstruction, we built the most complete model of the N-terminal pre-melastatin homology region. We also visualized several ligands that are bound by the protein and modeled how the human channel interacts with icilin. Analyses of pore helices showed that all available TRPM8 structures can be grouped into closed and desensitized states based on the register of pore helix S6 and the resulting positioning of particular amino acid residues at the channel constriction.

## INTRODUCTION

TRPM8 is a nonselective Ca^2+^-permeable transient receptor potential (TRP) channel. It belongs to the TRPM (melastatin) family that comprises eight members (TRPM1-8) that are involved in processing different stimuli, including temperature, taste, pressure, osmolality, and oxidative stress^1^. TRPM8 is a multimodal sensor of innocuous-to-noxious cold that is activated by several factors, such as temperature, cooling agents (e.g., menthol and icilin), voltage, pressure, and osmolality^2^. It is mainly expressed in sensitive primary afferent neurons that innervate cold-sensitive tissues, such as the skin, teeth, nasal mucosa, and tongue^3^. TRPM8 has attracted great interest because of therapeutic applications of its modulators that can be utilized for several pathological conditions, including neuropathic pain, irritable bowel syndrome, oropharyngeal dysphagia, chronic cough, and hypertension^4^. Importantly, TRPM8 is involved in various processes that are related to cancer progression, suggesting that its ligands can exert anticancer activity in tissues that express TRPM8, such as breast cancer, bladder cancer, esophageal cancer, lung cancer, skin cancer, pancreatic cancer, colon cancer, gastric cancer, and osteosarcoma^5^. Because of its importance, a significant number of TRPM8 modulators have been reported^6^. Nevertheless, to support further structure-based rational drug design, binding sites of this ion channel and the mechanism of its modulation by ligands, particularly for the human protein, need to be better explored^7^.

The first TRPM8 structures were solved by cryo-electron microscopy (EM) for proteins from birds (i.e., *Ficedula albicollis*, FaTRPM8^8,9^ and *Parus major*, PmTRPM8^10^). Both are highly homologous with the human TRPM8 channel, with sequence identity of 83% and 80%, respectively. The reported structures correspond to the apo and agonist-or antagonist-bound states. Notably, all avian structures are in the closed state, and binding of the ligand introduces only minor structural rearrangements. A much larger difference was observed for the structure that was solved in the presence of Ca^2+^ ions, which was defined as a desensitized state^10^. All avian TRPM8 structures show significant differences compared with the available mammalian structures of other TRPM channels (i.e., for TRPM4, TRPM5, and TRPM7), including conformation of the S4-S5 linker that connects transmembrane helices S4 and S5 in the apo state and the overall S5-S6 arrangement in Ca^2+^-bound structures^11^. Very recently, the first mammalian TRPM8 structures from mouse (MmTRPM8) were resolved^12^ in apo form, in a Ca^2+^-bound state, and in complex with icilin and Ca^2+^. The structures revealed that the mammalian TRPM8 channel, in contrast to most reported avian structures, has a canonical S4-S5 linker that resembles other mammalian TRPM proteins. MmTRPM8 contains a short selectivity filter that may account for its permeability to hydrated Ca^2+^. Ligand and Ca^2+^ binding induce very small conformational changes. Indeed, all resolved mouse structures share the same closed state that is very similar to the desensitized state of PmTRPM8^10^. Despite remarkable advancements in understanding TRPM8 that have been conferred by these avian and mouse structures that allowed TRPM8 activation and desensitization mechanisms to be proposed, several questions remain unanswered. Additional mammalian structures will further clarify the observed structural features and help interpret reported differences among TRP channels. Furthermore, no direct structural information is currently available for the human TRPM8 channel, and the N-terminal region that contains the pre-MHR and MHR1/2 domains is poorly resolved in all available structures.

To better understand structural features of the TRPM8 ion channel, we solved a cryo-EM structure of the human TRPM8 protein at 2.7 Å resolution. By combining a high-resolution global map with focused refinement maps of N-terminal domains, we were able to build an atomic model of Hs-TRPM8 that comprises residues 43-1104.

## RESULTS

### Cryo-EM structure of human TRPM8 channel

We determined a cryo-EM structure of apo human TRPM8 that was solubilized with lauryl maltose neopentyl glycol (LMNG) detergent. It corresponds to a closed state of the pore (Figure 1A, 1B, S1, and S2; Tables S1 and S2). At 2.7 Å resolution, it is one of the two highest-resolution structures of any TRPM8 protein that is currently available. Overall, human TRPM8 adopts a characteristic homotetrameric arrangement and is very similar to previously described TRPM8 structures^8-10,12^. The folded N-terminal part of the protein (residues ∼43-100), termed the pre-MHR, is followed by four MHRs (MHR1-4; Figure 1C, 1D). Together they comprise ∼630 residues and form the majority of the cytoplasmic part of the channel^13,14^. Melastatin homology regions are followed by a transmembrane channel domain (TMD) that is characteristic of all proteins from the TRP superfamily. It resembles TMDs that are present in TRPV1^15^ and TRPV2^16^ and is composed of a pre-S1 domain, a VSL domain, and the pore. The TMD is followed by the TRP helix that is involved in formation of the binding pocket for cooling agents^9,10,12^. The C-terminal domain (CTD) is composed of two helices (CTDH1/2) and a long coiled-coil element that extends toward the cytosolic part of the structure and plays key roles in stabilization of the tetrameric structure.

**Figure 1.**
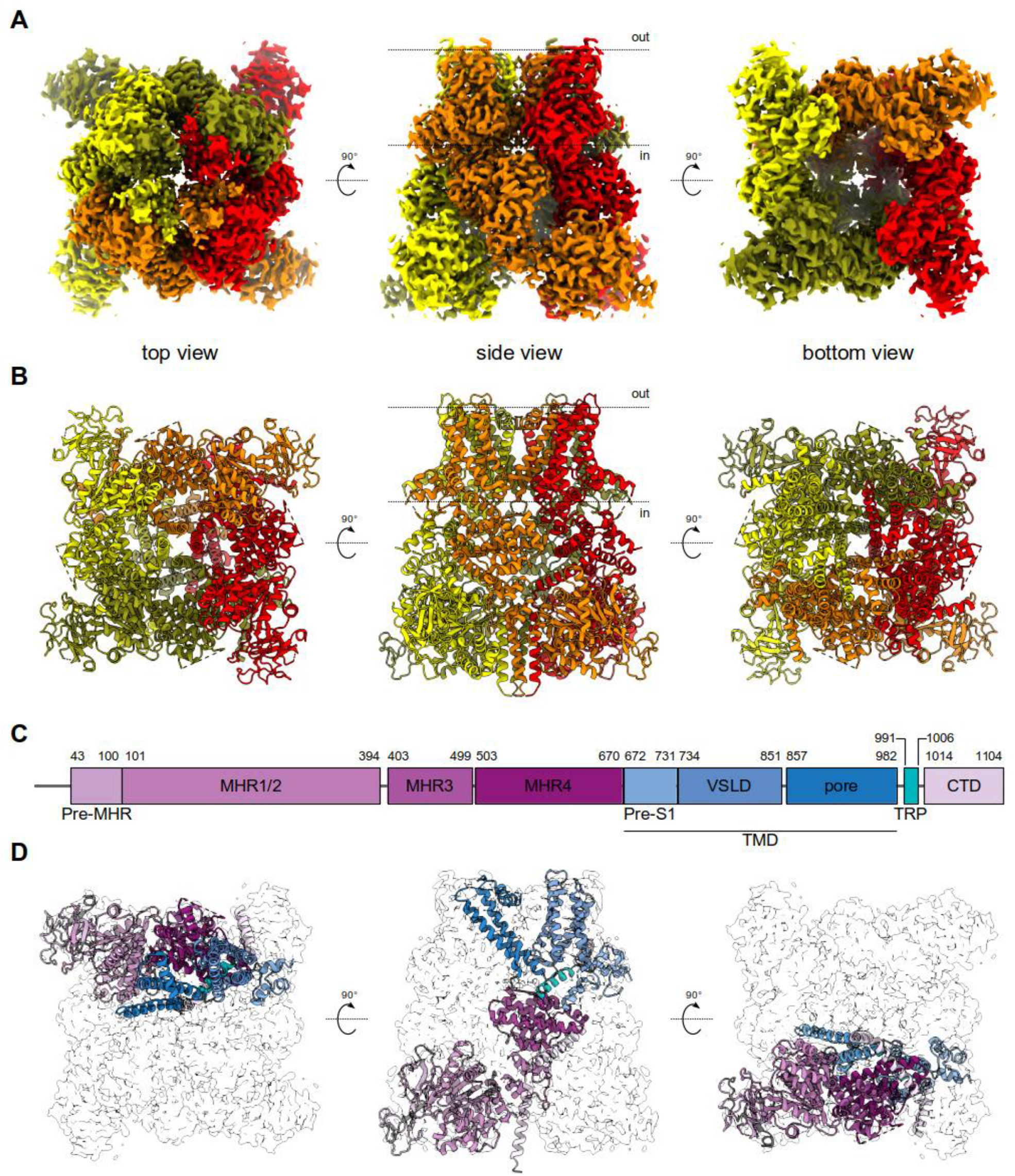
Overall structure and domain composition of human TRPM8 channel. (**A**) Cryo-electron microscopy electrostatic potential maps colored according to the channel subunits (olive, red, orange, and yellow; three views). (**B**) Atomic model of HsTRPM8 in cartoon representation, colored as in (A). (**C**) Domain composition of HsTRPM8. Residue numbers at the boundaries between particular elements are given. (**D**) Structure of a single subunit with domains colored as in (C). A contour of the cryo-EM map for the entire tetramer is shown in gray (three views). Dotted lines in (A) and (B) indicate the approximate boundaries of the cell membrane. See also Figure S1 and S2.

### The pore of HsTRPM8 is in a closed state

The transmembrane region of TRPM8 adopts a fold characteristic of proteins from the TRP family, with helices S4-S6 swapped between adjacent subunits of the tetramer. The pore is composed of helices S5, PH, and S6 (Figure 2A). The lower gate of the channel is formed by hydrophobic side chains of M978 and F979 (both shown in Figure 2A as sticks) that close the pore. The upper gate, formed by the selectivity filter followed by the outer pore loop (residues 914-951) that links the PH and S6 helices, is not fully resolved in our reconstruction. A large-scale structural comparison of all TRP channels^17^ suggested that selectivity filters of nonselective TRP channels have intrinsic flexibility. We speculate that the filter is stabilized in some specific conformations, thus allowing its visualization by cryo-EM.

**Figure 2.**
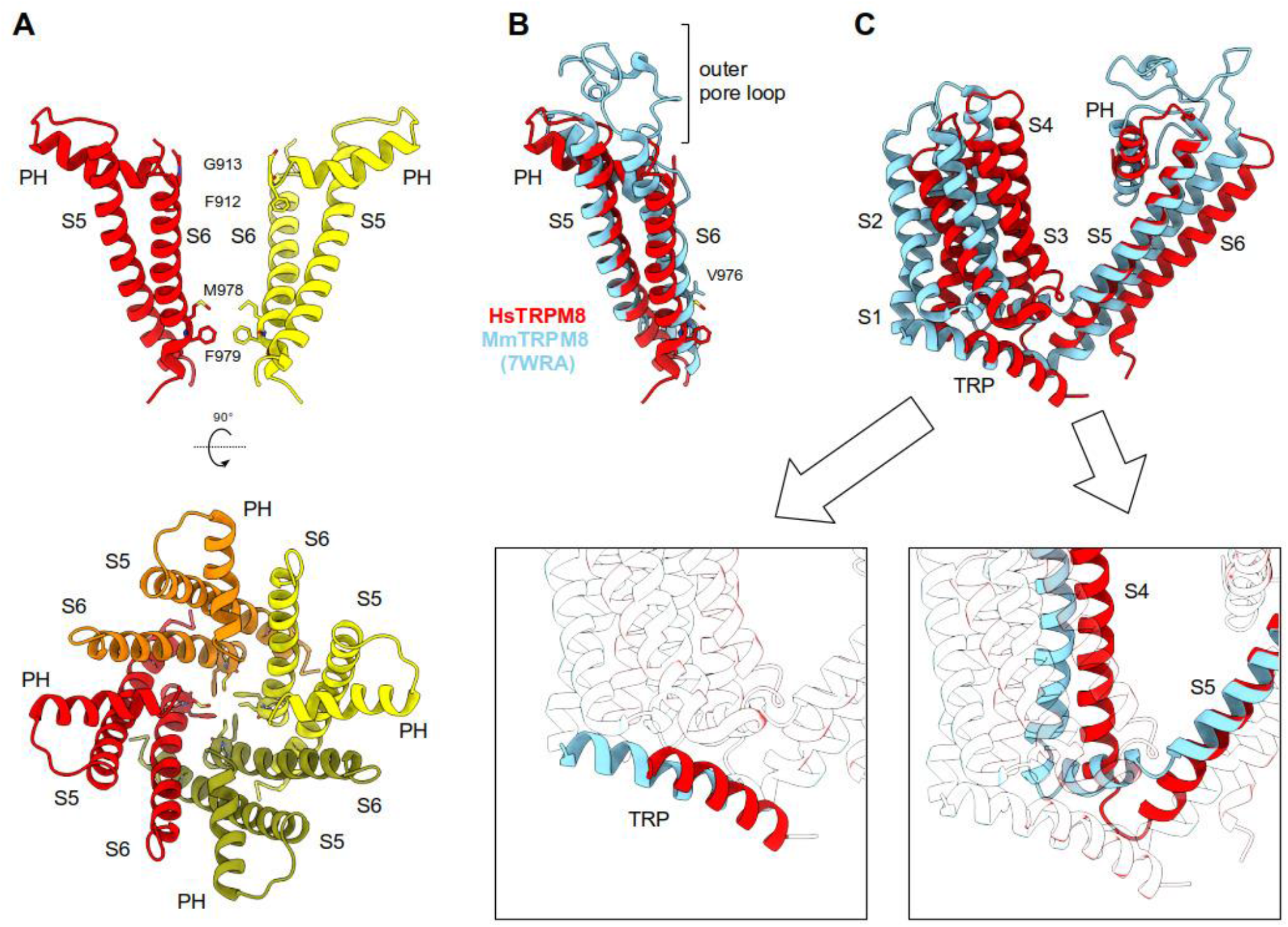
Configuration of the HsTRPM8 TMD in a closed state. (**A**) The pore of HsTRPM8, composed of helices S5, PH, and S6 (two views). Hydrophobic residues M978 and F979 closing the gate and the F912 and G913 from the selectivity filter are shown as sticks. In the upper panel, front and rear subunits were removed for clarity. (**B**) Comparison of the pore of HsTRPM8 (red) and MmTRPM8 (light blue) (PDB ID: 7WRA). (**C**) Comparison of TMDs of HsTRPM8 (red) and MmTRPM8 (light blue) after superimposition of the corresponding pore regions. Insets show details of differences in the conformation of TRP helix (left) and S4-S5 linker (right). See also Figure S3.

Conformation of the pore in our structure is different from the conformation that was observed in the recently published MmTRPM8 structures^12^, where the S6 helix is twisted, altering its register that positions a different residue (V976) to close the lower gate (Figure 2B). This rearrangement is accompanied by stabilization of the upper gate and selectivity filter. In contrast, in our reconstruction, only the first two amino acids from the selectivity filter (F912 and G913) are visible. Important differences are also observed in the VSL domain, composed of helices S1-S4. In the mouse structures, helices S1-S4 are shifted, and the conserved TRP helix is positioned more parallel to the membrane bilayer (Figure 2C, left inset). Finally, substantial differences are visible in the S4 and S5 helices (Figure 2C, right inset). In MmTRPM8, a canonical S4-S5 linker is formed, and the S4 helix adopts a 3_10_ helical conformation at residues 841-LRL-843. In our structure, the S4-S5 linker is parallel to the S5 helix, and the S4 helix is fully α-helical. Notably, the conformation that is observed for MmTRPM8 structures corresponds to the state that was previously defined as a desensitized state^2^.

We compared our model of HsTRPM8 with available structures of TRPM8 channels by superimposing the corresponding pore regions. Conformation of the pore was the same between the human structure and PmTRPM8 in apo form (PDB ID: 6O6A; Figure S3A). The same conformation was also observed for PmTRPM8 in complex with the antagonists TC-I 2014 (PDB ID: 6O72) and AMTB (PDB ID: 6O6R) and for FaTRPM8 in the apo state (PDB ID: 6BPQ) and in complex with the icilin analog WS12 (PDB ID: 6NR2). Notably, among TRPM8 proteins, a different pore conformation was observed only for the desensitized Ca^2+^-bound structure of PmTRPM8^10^ (PDB ID: 6O77) and for the recently published structures of MmTRPM8 (Figure S3B). Similar rearrangements, albeit to a lesser extent, can also be observed in the FaTRPM8 structure with Ca^2+^ ions and icilin bound at high occupancy (PDB ID: 6NR3). This conformational change is mostly driven by the binding of icilin by FaTRPM8, leading to a rearrangement in the S4 helix in which the α-helix is converted to a 3_10_ helix in the vicinity of R841^9^. These differences notwithstanding, all reported structures correspond to the closed state of the channel.

A similar analysis was performed for pore regions of other members of the TRPM family. Pore helices from MmTRPM4^18^ (PDB ID: 6BCJ) and MmTRPM7^19^ (PDB ID: 5ZX5) superimpose well with their MmTRPM8 counterparts (Figure S3C). Interestingly, the register of characteristic pore closing residues is also preserved for all mouse structures and is different from the register that was found in our HsTRPM8 structure and most published avian structures (Figure S3D). This implies that the MmTRPM4 and MmTRPM7 channels in the reported structures adopt a conformation that is similar to the desensitized state of PmTRPM8, which is different from the one that is found in our HsTRPM8 atomic model.

### Analysis of ligand binding pockets in TMD

The main ligand-binding pocket in TRPM8 proteins, termed the VSLD cavity, is formed by residues from transmembrane helices S1-S4^8^. Previous studies indicated that residues R842 and K856 contribute to the voltage dependence of TRPM8^20^, whereas R842, Y745, and Y1005 have been shown to interact with menthol^20,21^. This is in agreement with the cryo-EM structure of PmTRPM8 and FaTRPM8, which visualized several ligands (including antagonists) in this pocket^9,10,12^. We collected cryo-EM data for HsTRPM8 in complex with several ligands, but we could not observe a clear density in this cavity (data not shown). However, in our apo HsTRPM8 reconstruction, we observed additional densities that clustered in putative ligand binding pockets around the TMD (Figure 3A, blue). In agreement with previous reports^10^, we observed a discrete density in the VSLD cavity that partially overlaps with icilin in the FaTRPM8 structure^9^ (Figure 3B). Additional well-defined densities are also visible inside the channel between helices S5 and S6. We and others^10^ assigned these to an unidentified phospholipid tail (modeled as undecane) and two Na^+^ ions (Figure 3C). Finally, in the vicinity of the swapped helix S5 from one tetramer subunit and VSLD of the adjacent subunit, we observed clear densities that were assigned previously to cholesterol hemisuccinate (CHS), 3-SN-phosphatidylethanolamine (9PE), and another undecane molecule^10^. The quality of our EM map in this region allowed us to unambiguously assign them to two CHS molecules and a different type of phospholipid, phosphatidylcholine (POPC; Figure 3D). Because no specific ligands were added to our protein preparation, all these densities in our reconstruction likely correspond to molecules that co-purified with the protein or originate from chemicals that are used in protein purification and solubilization. Importantly, we did not observe any density in the putative Ca^2+^-binding pocket between the S2 and S3 helices that could be assigned to the divalent ion, which further confirms that our structure is not in the desensitized state.

**Figure 3.**
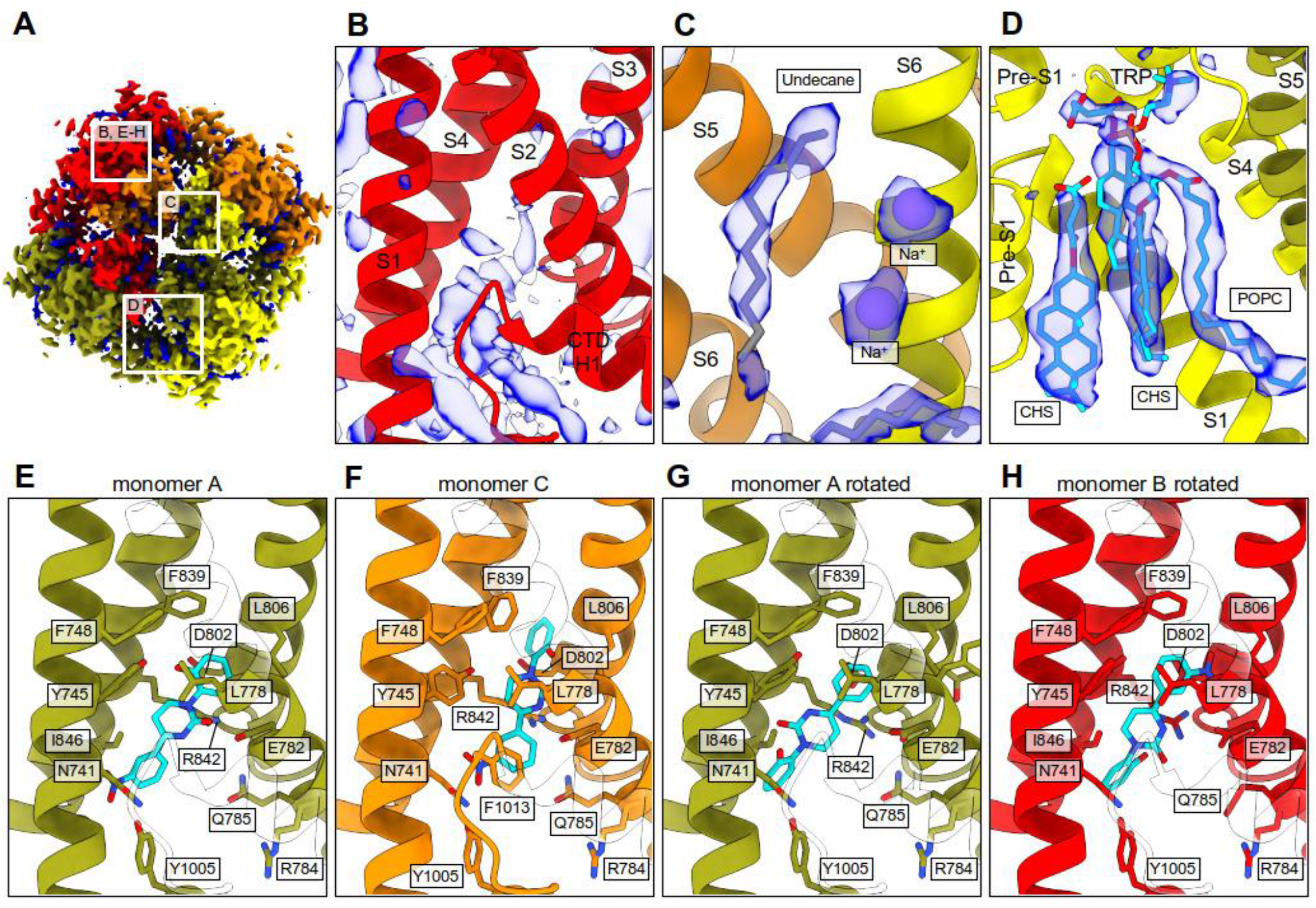
Ligand binding pockets in HsTRPM8 and modeling of icilin binding. (**A-D**) HsTRPM8 map (top view) with non-protein densities colored blue. (**B**) VLSD cavity with unidentified non-protein densities. (**C**) Densities between helices S5 and S6 assigned to undecane and Na^+^ ions. (**D**) Densities between helix S5 and VSLD assigned to two molecules of cholesterol hemisuccinate (CHS) and one phosphatidylcholine (POPC). (**E-H**) Modeling of icilin bound to VSLD cavity of HsTRPM8 in standard (E, F) and rotated (G, H) orientation. Predicted interacting residues are labeled and displayed as sticks. See also Figure S2E and Table S3.

Previous studies showed that binding to ligands does not significantly affect conformation of the pore, so we could use our structure to model a HsTRPM8 complex with icilin. By considering the two different poses that are assumed by icilin in the resolved structures^9,12^, two models were generated. The first model was obtained by superimposing the sensor module (S1-S4) of HsTRPM8 with avian FaTRPM8 in complex with icilin (PDB ID: 6NR3). The second model was obtained by manually rotating the icilin from the first complex by 180° to mimic the pose that is observed in the MmTRPM8 structure. The comparison of the complexes that were obtained for the four monomers of the HsTRPM8 tetramer in the first model reveals two different binding modes that mostly differ in the arrangement of the phenoxy ring (Figure 3E, 3F). Indeed, in all monomers, the dihydropyrimidinone ring approaches E782 and R842, and the nitrophenyl moiety interacts with N741 and Y1005. In contrast, only in one binding mode (represented by monomer A), the phenoxy group interacts with Y745. In the other mode (monomer C), it elicits clear π-π stacking with F839, whereas Y745 approaches the dihydropyrimidinone ring. The analysis of a set of representative scoring functions (Table S3) suggests that the complex of monomer C is more stable than the complex of monomer A, likely because of stronger hydrophobic contacts.

The analysis of rotated poses also revealed differences between the four subunits, mainly involving interactions that were formed by the E782/R842 dyad (Figure 3G, 3H). They point toward the nitrophenyl ring in monomer A. In monomer B, they assume a more central arrangement by which they contact all three rings. In both monomers, Y745 approaches the dihydropyrimidinone ring, whereas the phenoxy moiety contacts N741 and Y1005. The analysis of corresponding scores revealed that the complex of monomer B is more stable, mostly because of stronger polar interactions. Finally, the comparison of docking scores as computed for the original and rotated poses did not reveal significant differences, suggesting that both poses are similarly plausible within the explored binding site (Table S3).

### Structure of the Pre-MHR and MHR1/2 domains

In our initial reconstruction, fragments of the N-terminal region of the protein, spanning amino acids 1-400 (pre-MHR and MHR1/2 domains), were not clearly defined. Three-dimensional (3D) variability analysis in cryoSPARC suggested that this region is quite mobile, and its movements correlate with minor rearrangements of the whole pore structure (Movie S4). Interestingly, some of these movements appear to preserve two-fold symmetry rather than four-fold symmetry of the channel, which is in agreement with the reported TRPM2 structures that showed intermediate states with C2 symmetry^9^. These regions were also not well resolved in the other reported TRPM8 structures^8-10,12^. To improve quality of the reconstruction of this region we performed its focused refinement, followed by C4-symmetry expansion and subtraction of the signal from the rest of the TRPM8 molecule. This resulted in 3.2 Å reconstruction with quality that was sufficient for the *de novo* building of amino acids 43-140 of the protein chain that were not clearly visible in all previous reconstructions (Figure 4A).

**Figure 4.**
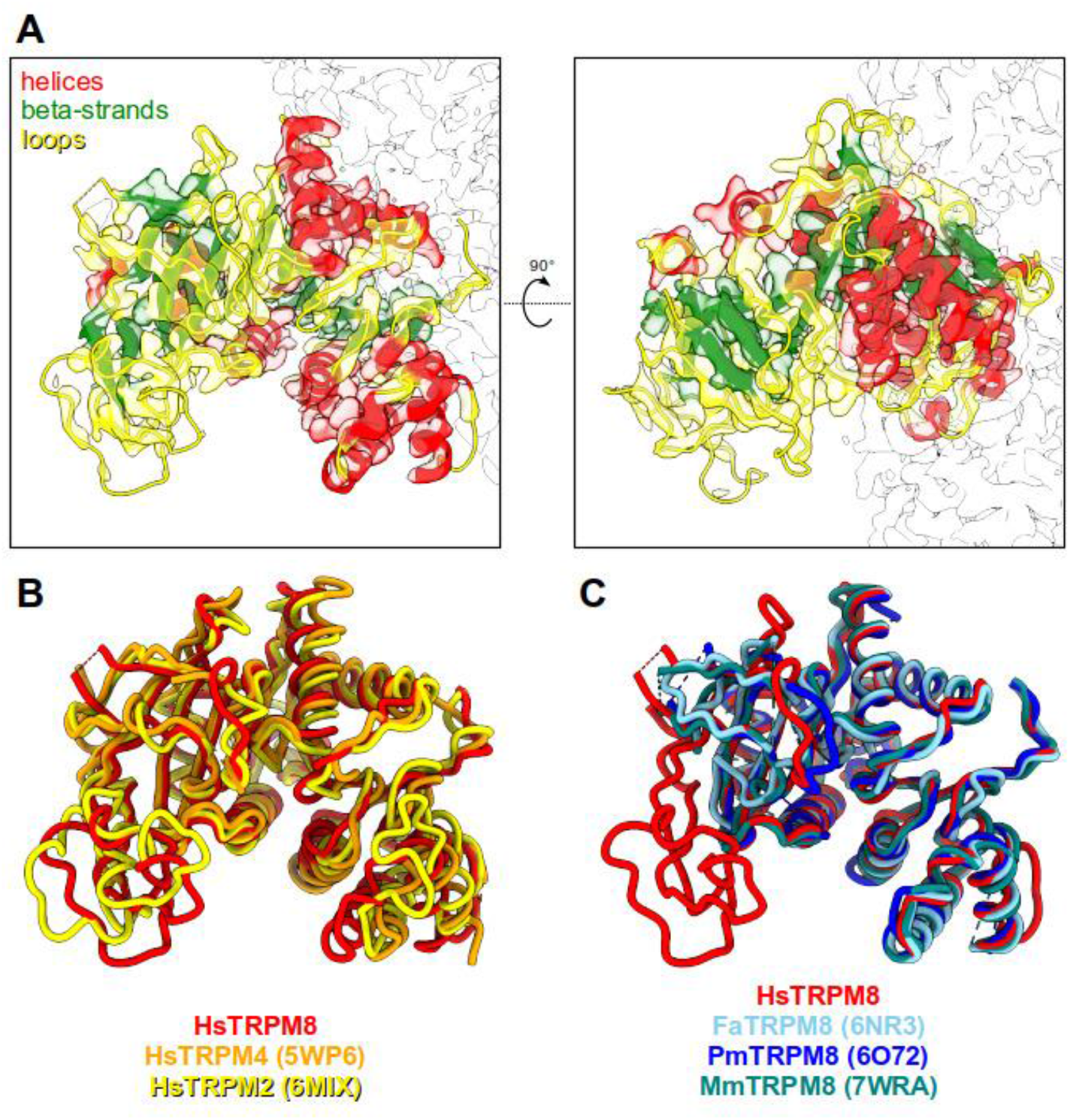
Structure of the pre-MHR and MHR1/2 domains of HsTRPM8. (**A**) Atomic structure of the pre-MHR and MHR1/2 from one subunit of the HsTRPM8 is shown in cartoon representation with helices in red, β-strands in green, and loops in yellow. A contour of the cryo-EM map for the entire tetramer is shown in gray (two views). (**B, C**) Comparison of pre-MHR from HsTRPM8 reported herein (red) with other human TRPM channels (B) and orthologous TRPM8 structures (C) (colored as in the figure keys).

The DALI search that was performed for this region showed that it adopts a canonical SLOG-like fold that is found in N-terminal regions of many other channels from the TRPM family, despite very low sequence identity (Figure 4B, < 30% identity for the shown structures). Interestingly, this region is involved in ligand binding in some TRP channels, suggesting its role in the sensing of external stimuli. In TRPM4, it is referred to as the N-terminal nucleotide-binding domain (NBD) and is responsible for binding adenosine triphosphate^18^ (ref). It also contains an additional binding pocket for the TRPM4 modulator decavanadate (DVT)^22^. The NBD is highly similar to the MHR1/2 domain in TRPM8, but the residues that are responsible for nucleotide binding and overall topology of the DVT-binding pocket are not conserved. We note that in our focused map, this region is better resolved (including a long loop that comprises residues 55-94) than in any other available TRPM8 structure (Figure 4C). Our model thus provides the most complete visualization of N-terminal portions of the TRPM8 channel to date.

## DISCUSSION

Recent years have brought a number of cryo-EM structures of TRPM8 channels^8-10,12^. Despite a broad spectrum of ligands that are used in these studies, all of them correspond to the closed state of the channel. The structural alignment of all available TRP channel structures^17^ revealed that the lower gate can be constricted at four sites (A, B, C, and D), spanning three helical turns of the S6 helix (Figure S3D). In most TRP structures, the pore is closed by a small and hydrophobic residue at site B (L/I/V), accompanied by a less conserved polar or hydrophobic residue from site C. All reported MmTRPM8 structures and the Ca^2+^-desensitized PmTRPM8 channel (PDB ID: 6O77) have the same S6 helix configuration. In contrast, the conformation of all other available TRMP8 structures (including the one reported herein) is different, with the register of S6 helix altered by half a turn. This forces different residues (“+2” according to the above register or “-2” for the FaTRPM8 protein that was solved with PIP2 and icilin at high occupancy [PDB ID: 6NR3]) to constrict the channel. In turn, in human and most avian structures, the pore is closed through hydrophobic interactions of side chains of F979 and M978 (site B), followed by T982 (site C). Overall, these differences in configuration of the S6 helix show flexibility of the lower gate and how it can respond to different stimuli.

The register shift in the S6 helix is accompanied by a number of conformational changes in the pore region. These include (*i*) a rigid-body tilt of the VSLD, (*ii*) the formation of a canonical S4–S5 linker, (*iii*) shifts of the S5, PH, and S6 helices, (*iv*) stabilization of the outer pore loop, (*v*) tilting of the TRP domain, and (*vi*) the introduction of a 3_10_-helix in S4 and π-helix in both the PH and S6 helices. All these rearrangements were collectively described as the transition from a closed to a desensitized state^2,10^. Our analyses imply that our HsTRPM8 structure resembles the closed state that is described for avian TRPM8 proteins, whereas the MmTRPM8 structures are indistinguishable from the desensitized state^10^. Another possibility is that HsTRPM8 and MmTRPM8 are simply two different conformations of a closed state. According to the maximum-likelihood analysis of voltage-dependent single-channel gating in cell-attached patches, there are at least five different closed states and two open states of the TRPM8 channel^23^. Thus, it is possible that both of the described conformations are simply different closed states of the TRPM8 channel. Further structural studies are required to fully describe the TRPM8 gating mechanism.

## MATERIALS AND METHODS

### Reagents and Chemicals

pFastBac Dual His6 MBP N10 TEV LIC cloning vector (5C) was purchased from Addgene (Addgene plasmid no. 30123; https://www.addgene.org/30123). All restriction enzymes, the pFastBac1 vector, and DH10Bac cells were purchased from Thermo Fisher Scientific. Sf9 cells were purchased from ATCC. Q5^®^ High-Fidelity DNA Polymerase, the Monarch^®^ DNA Gel Extraction Kit, and amylose resin were purchased from NEB (https://international.neb.com/). The DNA purification kit was purchased from Promega. The anti-6xHis antibodies were purchased from Abcam. All other chemicals and materials were purchased from Bio-Rad Laboratories, GE Healthcare, Sigma-Aldrich, or Roche Diagnostics, unless otherwise indicated.

### Plasmid Construction, Cloning, Expression, Purification of hsTRPM8

Synthetic cDNA that encoded full-length *Homo sapiens* TRPM8 (hsTRPM8, NCBI Reference Sequence NM_024080.5) was codon optimized for *S. frugiperda* and cloned into a pEX-K248 vector by Eurofins Genomics (https://eurofinsgenomics.eu/). cDNAs of 6xHis-MBP-TEV from the pFastBac Dual His6 MBP N10 TEV LIC cloning vector (5C) and hsTRPM8 from the pEX-K248 vector were polymerase chain reaction (PCR) amplified with Q5^®^ High-Fidelity DNA Polymerase, gel purified by the Monarch^®^ DNA Gel Extraction Kit, and subcloned into a pBastBac1 expression vector using the EcoRI restriction site. The final His-MBP-TEV-hsTRPM8 construct contained full-length unmodified sequence of human TRPM8. This construct was used for DH10Bac *E. coli* cell transformation with subsequent bacmid isolation by isopropanol precipitation from appropriately selected colonies. To prepare baculovirus, Sf9 insect cells were transfected by PCR-verified His-MBP-TEV-hsTRPM8-containing bacmids in a six-well plate according to the manufacturer’s protocol (Bac-to-Bac, Invitrogen). After approximately 6-7 days, the low-volume P1 generation virus that contained cell media was harvested, spun down, and stored at 4°C. The P1 generation cell medium was used to create the high-volume/high-titer P2 generation virus stock. To prepare P2, 25 ml of cells at a density of ∼2 million cells/ml were infected by P1 media at a 1:100-200 ratio and incubated for 6-7 days, harvested, spun down, stored at 4°C, and used to prepare large-scale His-MBP-TEV-hsTRPM8 expression.

Sf9 cells between the 5th and 20th passage were utilized for His-MBP-TEV-hsTRPM8 expression. Briefly, cells were kept in ESF 921 Insect Cell Culture Medium (Expression systems) that was supplemented with 10% (v/v) fetal bovine serum (Gibco) and 1% Antibiotic Antimycotic Solution (Gibco) in an orbital shaker incubator that was set at 130 rotations per minute (rpm) and 27°C. They were cultured in a sterile 2 L Erlenmeyer flask in a 400-500 ml volume to density approximately 1.5 million cells/ml. The culture was then infected by the addition of the P2 generation virus stock at a final ratio of 1:100-200 (v/v), incubated for 72 h, harvested, and frozen in liquid nitrogen. The pellet was stored at -80°C. The level of His-MBP-TEV-hsTRPM8 expression was checked by anti-6xHis antibodies.

The purification procedure was performed based on a previous report^10^ with some modifications. All steps were performed at 4°C. The Sf9 cell pellet from the 400-500 ml culture was suspended in 15 ml of ice-cold buffer (20 mM HEPES [pH 7.4], 150 mM NaCl, and 2 mM Na-EGTA [pH 7.5]) that was supplemented with a cOmplete Protease Inhibitor Cocktail Tablet and Phosphatase Inhibitor Tablet (Roche) and 1 mM phenylmethylsulfonyl fluoride. To solubilize and extract hsTRPM8, 5-6 mM LMNG (Anatrace) and 1-1.2 mM Cholesteryl Hemisuccinate Tris Salt (CHS, Anatrace) were added directly to the cell lysate, and the solution was stirred for 2 h. The insoluble material was removed by centrifugation at 30,000 rpm for 50 min at 4°C. Amylose resin (0.7-0.8 ml) was prepared by several washes with H_2_0 and a final wash with buffer and incubated with His-MBP-TEV-hsTRPM8-containing supernatant for 2-3 h. Amylose resin was transferred to a gravity-flow column and sequentially washed 5-6 times with a buffer (20 mM HEPES [pH 7.4], 150 mM NaCl, 2 mM Na-EGTA [pH 7.5], 0.15 mM LMNG, and 0.03 mM CHS). The protein was then eluted in the same buffer that was supplemented with 20 mM maltose (Sigma-Aldrich). The fractions were pooled and concentrated on a Millipore concentrator (WMCO 50 kDa), and the His-MBP-tag was cleaved by TEV protease for 16-24 h at 4°C (at a ratio of 1:10 [w/w] for TEV protease and the protein sample, respectively). Next, the TRPM8 sample was loaded onto a Superose 6 Increase size-exclusion column (GE Healthcare) that was equilibrated with SEC buffer (20 mM HEPES [pH 7.4], 150 mM NaCl, 2 mM Na-EGTA [pH 7.5], 0.025 mM LMNG, 0.005 mM CHS, and 100 µM Tris[2-carboxyethyl]phosphine (TCEP). Fractions that corresponded to the hsTRPM8 tetramers were checked by negative staining, pooled, and concentrated to ∼0.5 mg/ml using an Amicon centrifugal filter device (WMCO 100 kDa, Millipore) for cryo-EM analysis. We note that our structure was obtained using protein that was solubilized by LMNG. Structures of other TRPM proteins that were determined using different methods (e.g., crystal *vs*. cryo-EM or detergent/amphipol solubilization *vs*. nanodisc embedding) were essentially the same^17,24^. Therefore, the method of structure determination does not appear to affect the structure of these channels.

### Cryo-EM sample preparation and data collection

The HsTRPM8 tetramers that were solubilized by LMNG and concentrated to ∼0.5 mg/ml were applied to an UltrAuFoil 1.2/1.3 mesh 300 cryo-EM grid (Jena Bioscience, catalog no. X-201-Au300), glow-discharged from both sides. The sample was vitrified in liquid ethane using an FEI Vitrobot Mark IV (Thermo Fisher Scientific) at 4°C and 95% humidity with a 4 s blot time and 0 blot force. Data collection was performed with a Titan Krios G3i electron microscope (Thermo Fisher Scientific) that operated at 300 kV and was equipped with a BioQuantum energy filter (with 20 eV energy slit) and K3 camera (Gatan) at the SOLARIS National Synchrotron Radiation Centre (Krakow, Poland). Movies were collected with Aberration-free image shift (AFIS), with a nominal magnification of 105,000× (corresponding to a physical pixel size of 0.82 Å), 50 µm C2 aperture, and retracted objective aperture. The defocus range was set to -0.9 to -2.7 μm. The total dose (fractionated into 40 frames) was 41.42 e/Å^2^. The dose rate was 16.67 e/pixel/s, measured without sample (in vacuum).

### Cryo-EM data processing

A total of 7,918 movies were collected and processed with RELION-3.1^25^ and cryoSPARC v3.3.2^26^ (Figure S1). Raw movies were motion-corrected and binned twice using RELION’s implementation of MotionCor2 software^27^, and 1,090 micrographs with the total calculated motion larger than 100 pixels were discarded. The contrast transfer function (CTF) was fitted with CTFFIND-4.1^28^, and 5,714 micrographs with the maximum CTF resolution below 5 Å were selected for subsequent processing. A total of 1.24 million particles were picked with crYOLO^29^ and extracted with a binned pixel size of 3.28 Å. After two rounds of reference-free two-dimensional classification in cryoSPARC, 321,120 selected particles were re-imported into RELION with scripts from the University of California, San Francisco (UCSF) *pyem* package^30^ and re-extracted with a pixel size of 1.64 Å. Two additional rounds of two-dimensional classification in cryoSPARC further reduced the number of particles to 262,884. The selected particles were then subjected to 3D refinement (with the initial model created in cryoSPARC during the preliminary dataset processing) and again re-imported into RELION. Two rounds of 3D classification with local angular searches were then used to clean the particle stack. Three-dimensional refinement with an imposed C4 symmetry of 132,301 selected particles resulted in a 3.32 Å reconstruction. Refined particles were subjected to *Bayesian polishing* procedure (with un-binning to a pixel size of 0.82 Å), followed by CTF refinements, which improved resolution to 2.95 Å. After a second round of *Bayesian polishing*, particles were filtered with a third round of 3D classification. A total of 110,176 selected particles were imported into cryoSPARC for a final round of 3D refinement with imposed C4 symmetry, which resulted in a 2.65 Å reconstruction. This final map was sharpened locally with the *Local Filtering* tool in cryoSPARC with a B factor of -87 Å^2^.

For focused MHR1/2 refinement, the final set of particles was re-imported into RELION and subjected to *C4 Symmetry Expansion* procedure, which increased the number of aligned particles to 440,704. To facilitate the local alignment of a single MHR1/2 region, the signal from all other parts of the TRPM8 reconstruction (including three additional copies of MHR1/2) was removed from the images with a *Signal Subtraction* tool. After one additional round of 3D classification, 212,851 selected particles were imported into cryoSPARC for a final round of *Local Refinement*, which resulted in a 3.20 Å reconstruction. This focused map was sharpened locally with the *Local Filtering* tool in cryoSPARC with a B factor of -148 Å^2^.

The composite map was created in UCSF Chimera^31^ by combining the global map and four copies of focused maps with the *vop maximum* command. All reported resolutions were estimated from gold-standard masked Fourier shell correlation (FSC) curves at the 0.143 threshold. Data collection and processing statistics are present in Table S1.

### Cryo-EM map interpretation and model building

The composite map of HsTRPM8 was used for *de novo* model building in Coot^32^ with the aid of the orthologous structures PmTRPM8^10^ (PDB ID: 6O6A/6O72) and MmTRPM8^12^ (PDB ID: 7WRA) and the AlphaFold^33^ model for the N-terminal part of MHR1/2. Multiple rounds of real-space refinement in Phenix^34^ and model building in Coot resulted in an almost complete structure of HsTRPM8 that comprised residues 43-1104, with several loops and disordered regions missing (residues 229-234, 534-556, 717-721, 914-951, 985-989, 1031-1048).

Local resolution of the HsTRPM8 map was the highest for the central part of the molecule (i.e., the pore and MHR3/4). Local resolution was the lowest for extracellular loops and C-terminal coiled-coil elements (Figure S2A, S2B). Map quality was sufficient to build most side chains in well-structured pore regions and unambiguously trace the protein backbone and some of the side chains in the lower-resolution MHR1/2 (Figure S2C, S2D). In agreement with previous reports^10^, we observed additional densities in the transmembrane region that we modeled as two CHS molecules and one POPC (Figure 3D; Figure S2E). Additionally, in the center of the pore, near the extracellular part, we observed clear densities that we modeled as two Na^+^ ions (the cation at the highest concentration in the sample) and a single undecane molecule^10,12^ (presumably part of an unidentified phospholipid tail; Figure 3C).

All figures were prepared in UCSF ChimeraX^35^. Refinement and validation statistics are present in Table S2.

### Modeling of icilin binding

The icilin complexes were generated by superimposing the hTRPM8 structure with the avian faTRPM8 6NR3 structure. Superimposition was guided by backbone atoms of the sensor module only (S1-S4 helices) and was performed using the Vega suite of programs^36^. By considering the different arrangements as detected for icilin in the resolved structures, an alternative structure was generated by rotating 180° the previously determined icilin poses. These obtained complexes were then minimized using NAMD^37^ by keeping fixed all atoms outside an 8 Å radius sphere around the icilin. Additionally, all backbone atoms were kept fixed to preserve the experimental folding.

---

**Figure S1.**
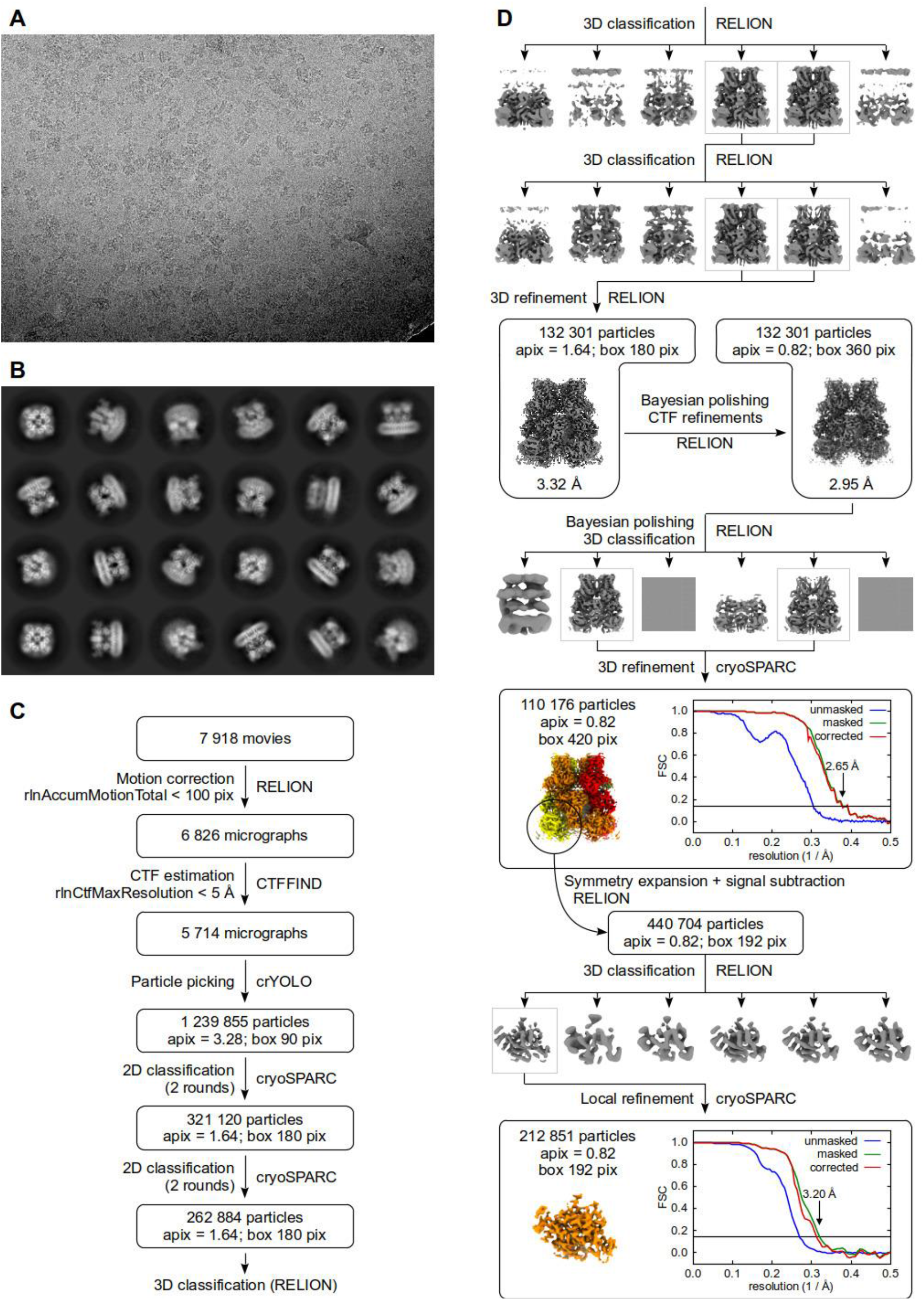
Cryo-electron microscopy data processing, related to Figure 1. (**A**) Representative cryo-EM micrograph. (**B**) Selected class averages after last round of two-dimensional classification. (**C**) Initial processing steps and particle curation. (**D**) Three-dimensional reconstruction pipeline. For the final reconstructions, gold-standard Fourier shell correlation (FSC) curves between two half maps are presented. The horizontal line represents a value of 0.143. Final maps are colored as in Figure 1.

**Figure S2.**
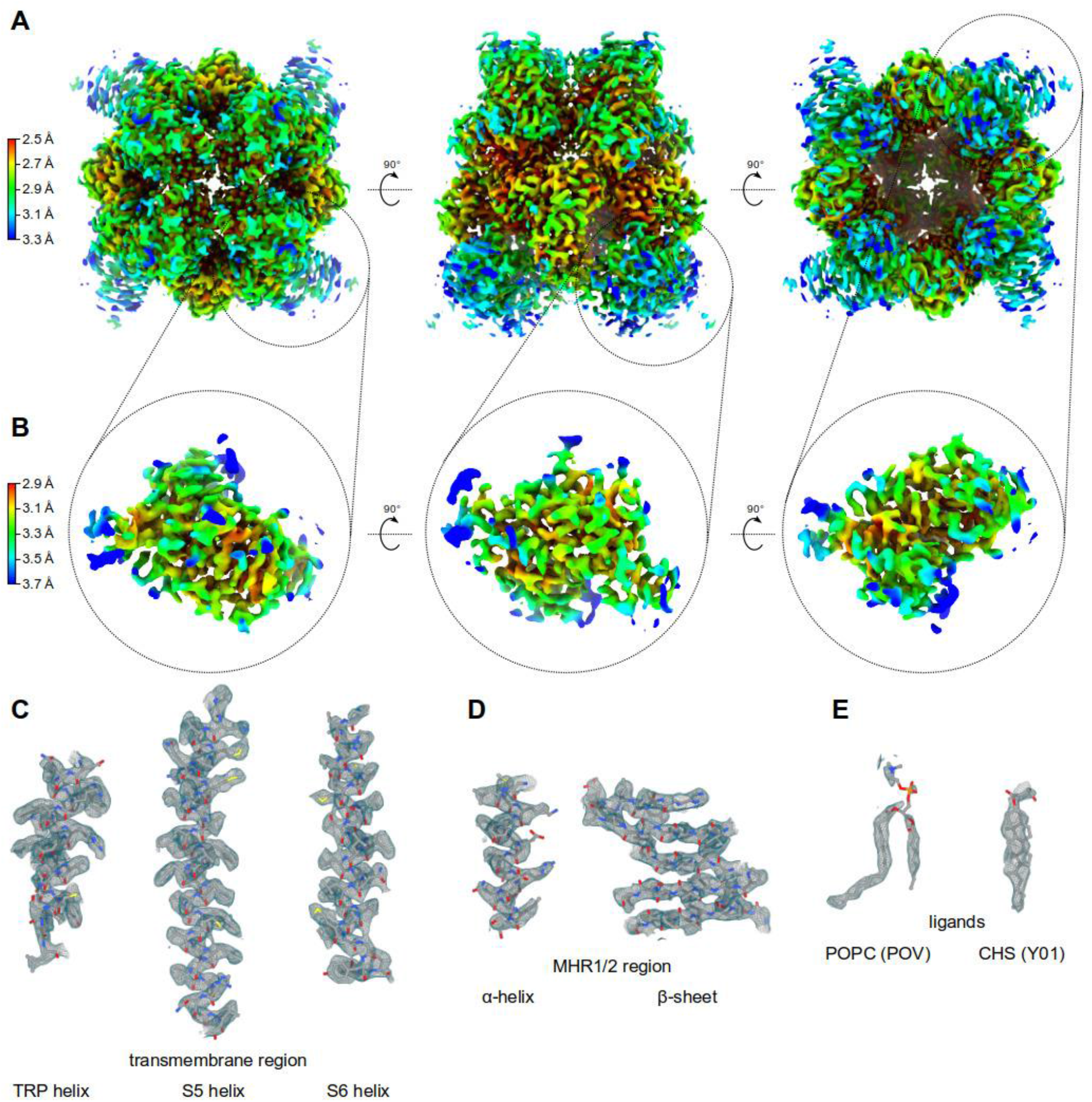
Quality of cryo-EM maps, related to Figure 1, Figure 3, and Figure 4. (**A, B**) Local resolution calculated from half maps in cryoSPARC for (**A**) consensus TRPM8 refinement and (**B**) focused MHR1/2 refinement. (**C-E**) Quality of cryo-EM map, showing selected secondary structures from the pore region (**C**) and the MHR1/2 domain (**D**). (**E**) Electron microscopy potential density for the selected modeled ligands: phosphatidylcholine (POPC) and cholesterol hemisuccinate (CHS).

**Figure S3.**
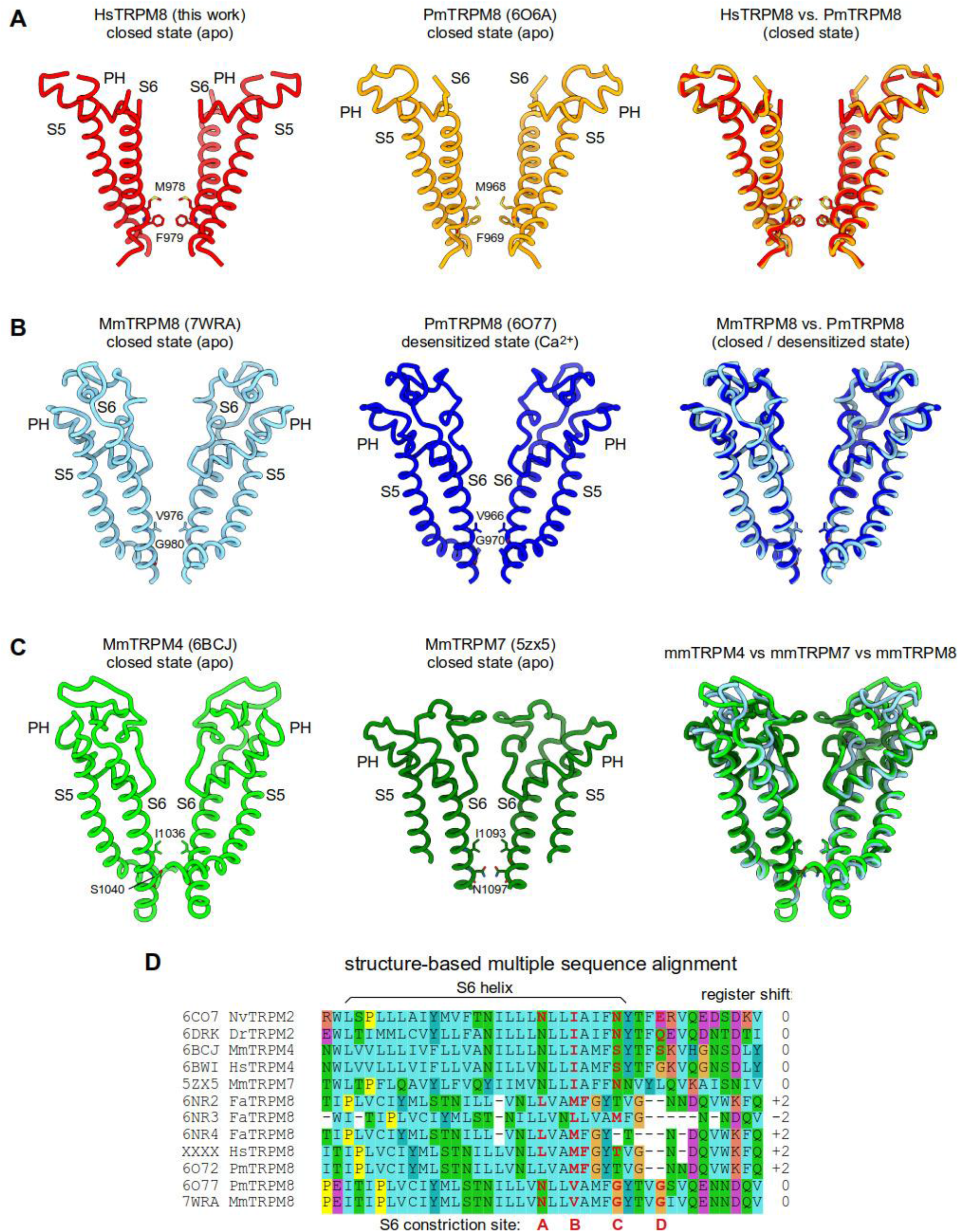
Comparison of the pore region of selected TRPM channels, related to Figure 2. (**A**) TRPM8 structures in closed state: HsTRPM8 (this work) and PmTRPM8 (PDB ID: 6O6A). (**B**) MmTRPM8 structure (PDB ID: 7WRA) aligned with desensitized PmTRPM8 structure (PDB ID: 7O77). (**C**) MmTRPM8 (PDB ID: 7WRA) aligned with MmTRPM4 (PDB ID: 6BCJ) and MmTRPM7 (PDB ID: 5ZX5). Front and rear subunits were removed for clarity. (**D**) Structure-based sequence alignment of S6 helices from selected TRPM channels. See also Figure 4 in Huffer et al., 2020^17^

**Table S1.**
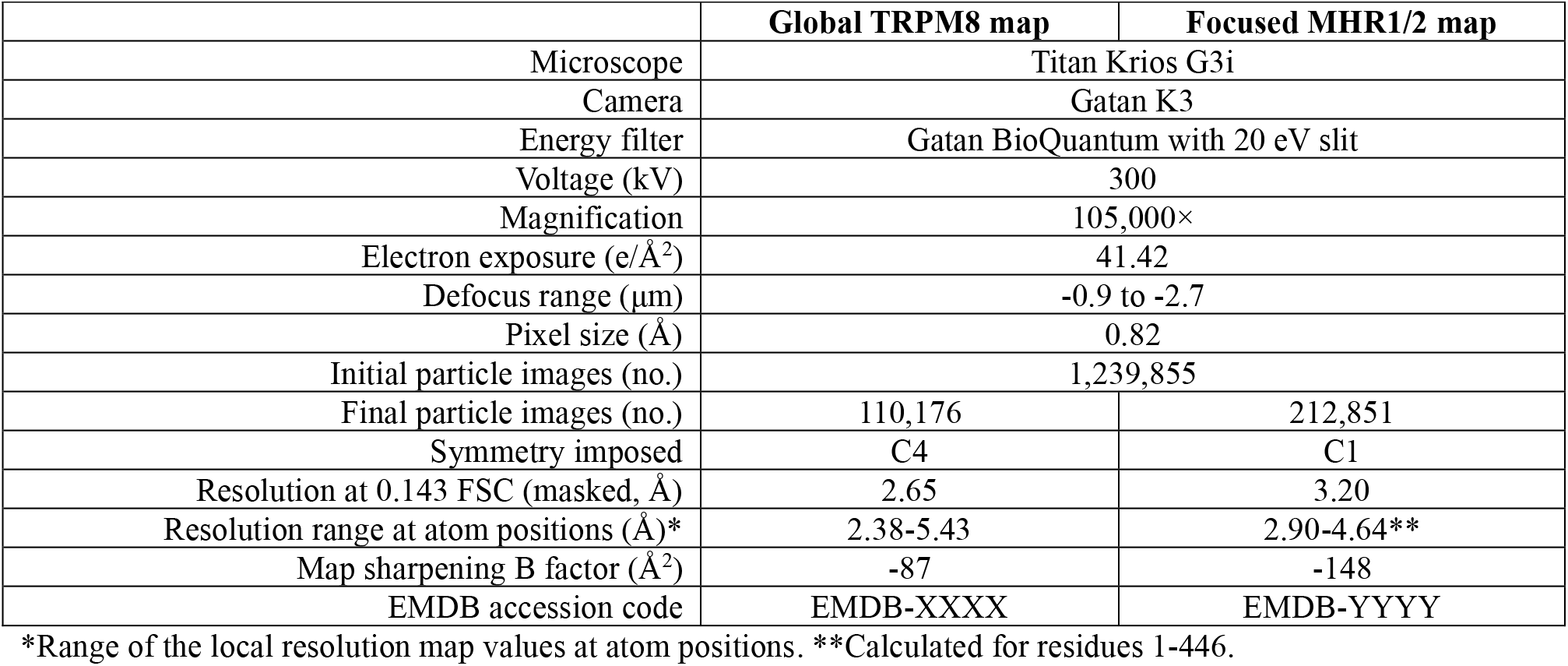
Data collection and processing.

**Table S2.**
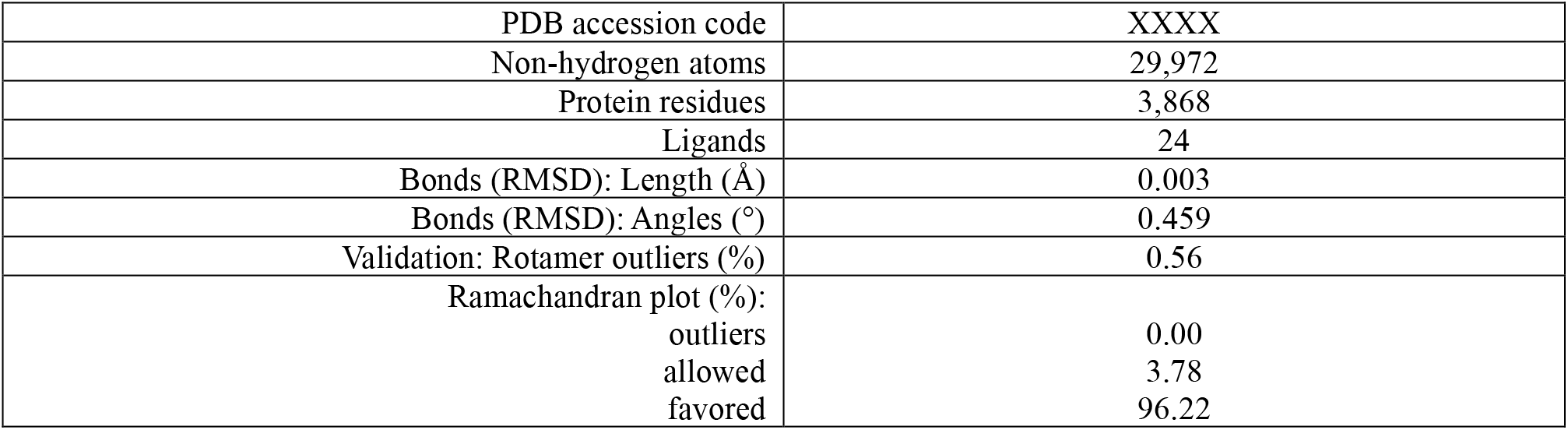
Refinement and validation statistics.

**Table S3.** Docking scores for modelling of icilin.

|  | **Xscore** | **ChemPLP** | **Elec_DD** | **No. contacts** | **CHARM6-12** |
| --- | --- | --- | --- | --- | --- |
| Original pose |  |  |  |  |  |
| A | -9.05 | -59.14 | -2.19 | 6 | -41.99 |
| B | -9.05 | -58.92 | -2.20 | 6 | -41.99 |
| C | -9.05 | -86.45 | -4.39 | 11 | -44.77 |
| D | -9.23 | -72.53 | -3.72 | 11 | -40.49 |
| Mean | -9.10 | -69.26 | -3.12 | 8.5 | -42.31 |
| Rotated pose |  |  |  |  |  |
| A | -9.23 | -49.84 | -1.51 | 8 | -40.90 |
| B | -9.42 | -81.32 | -3.82 | 7 | -40.41 |
| C | -9.31 | -60.73 | -3.48 | 8 | -40.06 |
| D | -9.17 | -80.75 | -1.93 | 7 | -38.94 |
| Mean | -9.28 | -68.16 | -2.68 | 7.5 | -40.08 |

The XScore and ChemPLP scoring functions encode for overall stability of the generated poses. The Elec\_DD score focuses on electrostatic interactions as computed by applying a distance dielectric function. The CHARM6-12 score corresponds to the non-bond Lennard Jones interaction energy using CHARMM parameters. “No. contacts” refers to residues within a 3 Å radius sphere around the ligand. Apart from “No. Contacts,” which is dimensionless, all other scores are expressed in kcal/mol.

